# Quantitative coupling of cell volume and membrane tension during osmotic shocks

**DOI:** 10.1101/2021.01.22.427801

**Authors:** Chloé Roffay, Guillaume Molinard, Kyoohyun Kim, Victoria Barbarassa, Marta Urbanska, Vincent Mercier, José García-Calvo, Stefan Matile, Jochen Guck, Martin Lenz, Aurélien Roux

## Abstract

During osmotic changes of their environment, cells actively regulate their volume and plasma membrane tension that can passively change through osmosis. How tension and volume are coupled during osmotic adaptation remains unknown, as a quantitative characterization is lacking. Here, we performed dynamic membrane tension and cell volume measurements during osmotic shocks. During the first few seconds following the shock, cell volume varied to equilibrate osmotic pressures inside and outside the cell, and membrane tension dynamically followed these changes. A theoretical model based on the passive, reversible unfolding of the membrane as it detaches from the actin cortex during volume increase, quantitatively describes our data. After the initial response, tension and volume recovered from hypoosmotic shocks but not from hyperosmotic shocks. During these asymmetric recoveries, tension and volume remained coupled. Pharmacological disruption of the cytoskeleton and functional inhibition of ion channels and mTOR all affected tension and volume responses, proving that a passive mechanism is necessary and critical for the cell to adapt fast. The coupling between them was, nonetheless, maintained for a few exceptions suggesting that volume and tension regulations are independent from the regulation of their coupling.

## Introduction

Lipid membranes are self-assembled viscoelastic bilayers separating cells and their organelles from their environment. They are easy to bend but resistant to stretching: their lysis tension - the tension at which they break - is high, in the range of a few mN/m^1, 2^. This high value protects cells against lysis upon processes that stretch the cell membrane. Plasma membrane tension arises from the combined contributions of osmotic pressure, in-plane tension and cytoskeletal forces^3, 4^. The cytoskeleton is intimately linked to all processes regulating membrane tension, in particular cell volume regulation^5^. For example, hypotonic shocks are not only responsible for increasing membrane tension but also induce the degradation of vimentin^6^ and a reorganization of actin filaments^7, 8^, without affecting microtubules^6^. It has been proposed that the cytoskeleton regulates membrane tension by setting its value through active force generation, and by establishing a membrane reservoir that buffers acute changes in tension^9^. This membrane reservoir is stored around protruding actin-based structures such as ruffles, filopodia and microvilli^10^. Cell volume regulation during osmotic changes involves mechano-sensitive ion channels^2, 11^ regulated by membrane tension^12, 13^. How ion channel activity is coupled to the cytoskeleton is under debate^14^. The channels comprise volume-regulated anion channels (VRACs), sodium-hydrogen antiporters (NHEs) and Na-K-Cl cotransporters (NKCC1). VRACs are activated by hypotonic stress^15, 16^ and are unique in transporting small organic osmolytes – in particular taurine - in addition to anions^17, 18^. NHEs inhibition prevents regulatory volume increases of cells^19, 20^. Cells have evolved to respond to changes in membrane tension so as to control their impact on many processes essential to cell life^21^. The genetic response to an osmotic stress has been studied extensively. This pathway partly consists of activating genes involved in the synthesis or degradation of osmo-protectant molecules (such as glycerol in yeast, and amino acids in mammalian cells), and their subsequent secretion^19, 22–26^. However, the genetic response minute timescale cannot account for the cell’s immediate resistance to stretch^19, 23^. The master regulator of plasma membrane tension is probably Target of Rapamycin Complex 2 (TORC2)^27^ and its mammalian homologous mTORC2. Indeed, TORC2 signaling increases instantaneously upon membrane tension increase^28^ as well as mTORC2 activity^29^, and decreases upon tension loss^26^. TORC2 regulates endocytosis through membrane tension^30^, but also actin polymerization^31^. Despite its undeniable importance, the mechanisms driving the regulation of membrane tension during osmotic shocks in relation to cell volume changes are still not understood. Qualitatively, membrane tension has been reported to decrease in response to hypertonic shocks^32, 33^, while studies have reported that it either stays constant^33, 34^ or increases^32, 35, 36^ upon hypotonic shock. The relation between osmolarity and cell volume change is captured by the Ponder/Boyle/Vant’Hoff (PBVH) relation whereby the cell shrinks until the osmotic pressure of its contents matches that of the extracellular medium^37^. This relation involves an osmotically inactive volume (OIV) which represents the minimum volume occupied by tightly packed cellular constituents at very high hypertonicity^38, 39^. In addition, while the PBVH relation describes the changes in cell volume in response to an osmotic shock, the response of the membrane tension to such shocks has never been quantitatively studied. In this study, we elucidate quantitatively the coupling between cell volume and membrane tension in single cells during osmotic shocks using time-resolved techniques.

## Results

We exposed HeLa Kyoto cells to step-wise osmotic shocks (Fig 1a, see Methods). A few seconds after a hypotonic shock with 75% water, cell volume peaked at approx. 2.4 times the initial volume. The volume subsequently recovered, but only partially, leaving a 15% volume increase after 10 min (Fig 1b). Weaker dilutions (50% and 25%) led to lower peaks and faster recovery (Fig 1b). Conversely, hypertonic shocks led to a rapid volume decrease within seconds, followed by a 10 min plateau. In the most extreme hypertonic conditions (3500 mOsm), cell volume decreased by up to 90% (Supp Fig 1a-b). Before the osmotic shocks, cell volume distributions were broad (Fig 1c, Supp Fig 1c-d) while relative cell volume changes were highly reproducible and essentially due to cytoplasmic volume changes (Supp Fig 1e-g). To assess the robustness of the recovery dynamics, we performed cell volume measurements using a different cell type (HL-60/S4) using real-time deformability cytometry (RT-DC) – a high-throughput technique allowing for rapid characterization of thousands of cells^40^ (Supp Fig 2a). After applying the osmotic shocks to HL-60/S4 cells before loading them into RT-DC, we confirmed our previous observation: cell volume in a hypotonic medium peaked and recovered while cell volume in a hypertonic medium abruptly and stably decreased (Supp Fig 2b-c). Our results show that the volume changes associated with osmotic shocks are rapid, and show a recovery for hypotonic shocks which is absent for hypertonic shocks and further demonstrate that cell volume can recover from hypotonic, but not hypertonic, shocks.

**Figure 1.**
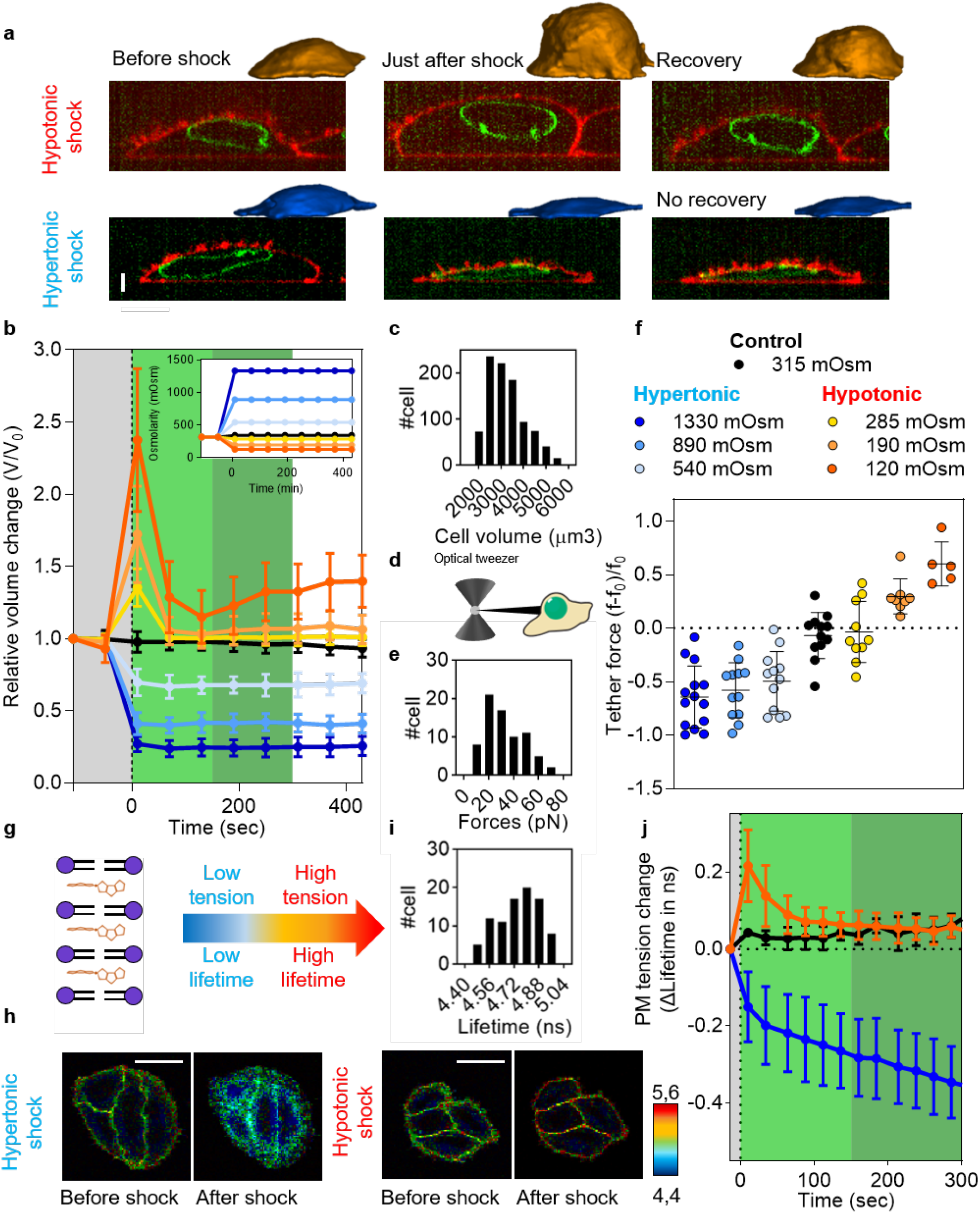
Osmotic shocks affect cell volume and membrane tension. **a**, 3D reconstruction of cell volume using Limeseg (top hypo, bottom hyper). **b**, Averaged cell volume dynamics under osmotic shock (grey: before shocks; light green: short-term response; dark green: long-term response). Insert: osmolarities (mOsm) of cell media with time for the different shocks. **c**, Cell volume distribution in isotonic medium before osmotic shocks. **d**, Tether forces pulled out from cells can be measured with optical tweezers. **e**, Tether force distribution in isotonic medium before osmotic shocks. **f**, Relative change of tether force immediatly after osmotic shocks (averaged over 10 sec) for different osmotic shocks. **g**, Fluorescence lifetime of the Flipper-TR probe reports membrane tension changes. **h**, FLIM images of Flipper-TR lifetime values (colorscale) of cells subjected to osmotic shocks. **i**, Distribution of the cell average Flipper-TR lifetimes in isotonic medium before osmotic shocks. **j**, Dynamics of the change of tension as measured by Flipper-TR lifetime (grey: before shock; light green: short-term response; dark green: long-term response).

We measured the dynamical changes of membrane tension after osmotic shocks. When membrane tubes are pulled from the cell membrane using beads held with optical tweezers (Fig 1d), the force required to hold the tube is a direct measurement of the membrane tension^41^. The distribution of tube forces before osmotic shock *f*_0_ is broad: 27± 18pN (Fig 1e). As seen for volume variations, changes in the tube force upon osmotic shocks were almost instantaneous (Supp Fig 2d). Interestingly, changes of the tube force (*f* – *f*_0_)/*f*_0_ averaged over 10 seconds immediately following the shock were proportional to the intensity of the shocks (Fig 1f, Supp Fig 2e-f). To follow the dynamics of tension in real-time, we used the molecular probe Flipper-TR (fluorescent lipid tension reporter, or FliptR^®^) that reports changes of membrane tension through changes of its fluorescence lifetime (Fig 1g-h)^26, 30, 33, 42–16^. Consistently with tube pulling experiments, the lifetime distribution of Flipper-TR in cells membrane before shock was broad: 4,76 ± 0,15 ns (Fig 1i), and the lifetime changed within seconds after shock (Fig 1h–1j). We observed an asymmetry in lifetime measurement during recovery phase similar to the one observed for volume measurements: membrane tension peaked and recovered within seconds after hypotonic shock while it decreased within seconds after hypertonic shock, and continued decreasing for the duration of the experiment, although at a lower rate (Fig 1j). Our results show that membrane tension variations after osmotic shocks qualitatively follow cell volume changes.

To quantitatively capture the relationship between the osmotic pressure of the cell and its volume (Fig 2a, Supp Fig 3a-c), we used the PBVH equation of state^37, 39, 47^

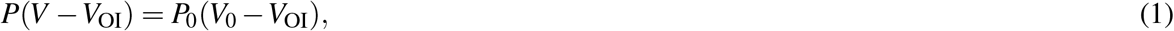

where *P* is the osmotic pressure of the intracellular medium, *V* the cell volume and *P*_0_ and V_0_ are values of P and V under isotonic conditions. Equation (1) assimilates the contents of the cell to a solution of particles with steric repulsions and otherwise negligible interactions, with the sum of the particles excluded volumes equal to VOI. The cell volume thus cannot be compressed below the ‘osmotically inactive volume’ *V*_OI_. Equation (1) is in excellent agreement with both our hypertonic and hypotonic data (Fig 2a), when the volume is estimated at the time of hypotonic peak (10s after the shock, shown in Fig 1b, Supp Fig 1b). A single-parameter fit yields *V*_OI_ = 300 *μm*^3^ equal to about 10% of the initial cell volume, smaller than previous estimates^48^. Interestingly, during and after the recovery phase (t>10s after shock) volume values diverge from this linear relation only in the hypotonic conditions, reflecting the asymmetry of recovery (Fig 2a). This explains why previous volume measurements after hypotonic shocks were not in agreement with the PBVH relation, probably because they were performed too late after the shock, at a time when cells had already recovered.

**Figure 2.**
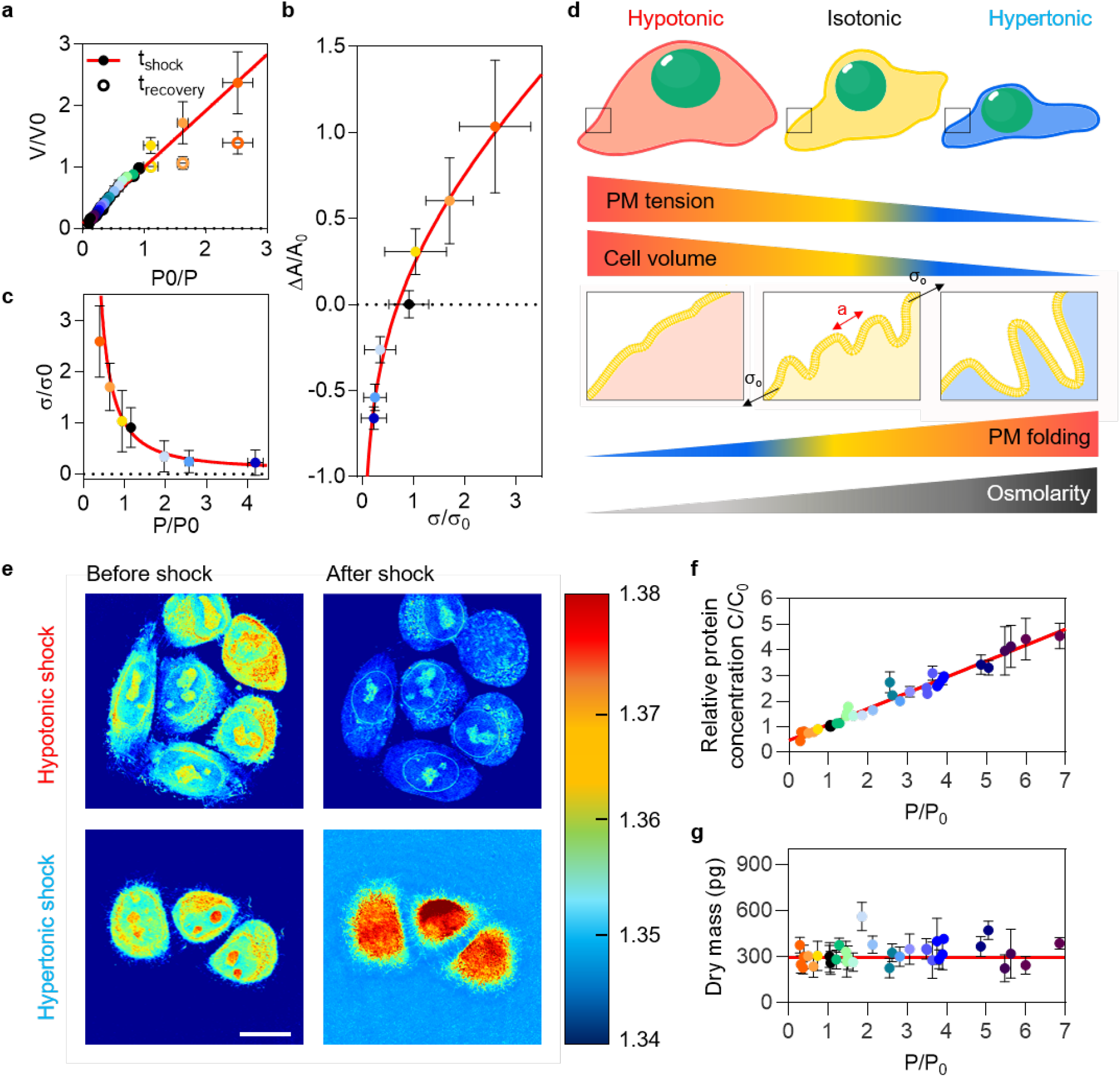
Quantitative coupling of cell membrane tension to osmotic shocks. **a**, Scheme describing the theory. **b**, Normalized volume changes (*V/V*_0_) as a function of osmotic pressure ratio (*P*_0_/*P*) just after osmotic shocks (full circle) and 8 min after the osmotic shock (empty circle, recovery phase) compared to the prediction of Eq. (1) (red line). **c**, Relative changes of membrane area (Δ*A/A*_0_) versus relative changes of membrane tension (Δσ/σ_0_) compared to the prediction of Eq. (2) (red line). **d**, Normalized tension (σ/σ_0_) versus normalized pressure (*P/P*_0_) and prediction obtained by combining Eq. (1) and Eq. (2). **e**, Refractive index images of cells under osmotic shocks. **f**, Protein concentration changes (*C/C*_0_) according to pressure applied (*P/P*_0_). **g**, Calculated dry mass of cells versus normalized pressure (*P/P*_0_)

As the volume of the cell changes, so does the tension and area of its membrane (Fig 1f–1j, Supp Fig 2d-f). To compute the relation between these quantities, we reasoned that the cell membrane is not a perfectly flat, but is instead heterogeneously folded because of protrusions and buds induced by proteins and cytoskeletal structures. The ruffled structure of the membrane would thus provide a large area buffer^9^. Increases in membrane tension unfold these ruffles, releasing more membrane area and allowing the cell to expand. To quantitatively model this effect (see theoretical supplement), we described the membrane energy by a modified Helfrich-Hamiltonian where the spontaneous curvature is randomly distributed according to a Gaussian distribution with exponential correlations. Its statistical properties were described by two parameters: the typical lateral ruffle size *a* (the correlation length of the noise), and the typical magnitude of mean curvature C (its strength). Under these assumptions, the relative change of membrane area during the shock is given by

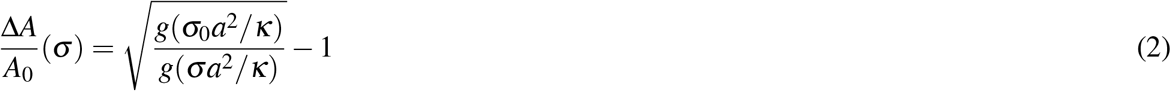

where σ_0_ and σ respectively denote the tension of the membrane in the initial isotonic state and in the final state, *κ* is its bending rigidity and the function *g* is given by

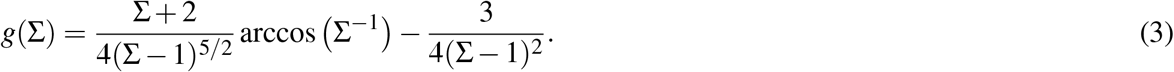

We find that Eq. (2) agrees very well with our experimental measurements (Fig 2b). The fit gives σ_0_ = 1.2 × 10^−4^N · m^−1^ and a ruffle size *a* = 37 nm, which are in good agreement with typical cell membrane tensions^49^ and the size of the smallest membrane invaginations such as caveolae and endocytic buds^50^. Finally, combining Eqs. (1), (2) and (3) yields a prediction for the dependence of the membrane tension on osmotic pressure, which is in good agreement with our data (Fig 2c). These results strongly support the notion that the short-term response of cell volume and membrane tension are predominantly mechanical and thermodynamic, and consists in a passive equalization of the inner and outer osmotic pressures accompanied by an unfolding of membrane ruffles (Fig 2d).

Our verification of the PBVH relation yields two surprising results: first, we find that it holds for a very large range of osmotic pressures in HeLa cells, larger than previously tested. Second, the *V*_OI_ represents a smaller proportion of *V*_0_ (only 10%, as compared to values between 7% and 50% in other studies). To better understand these results, we used optical diffraction tomography, a 3D tomographic label-free technique, to measure the cells refractive index (RI)^51, 52^ hence giving a direct access to changes of mass and concentration in cells subjected to osmotic shocks (Fig 2e). Cells in isotonic conditions had an average RI = 1.37± 0.01. A few seconds after 1M sucrose shock, cells had an increase in RI to 1.42± 0.01, while under 75% water shock conditions the RI decreased to 1.35± 0.01. RI increases linearly with increasing protein concentration^53^. In our experiment, the RI of single cells changed proportionally to the applied osmotic pressure (Supp Fig 3d). This implies that the protein concentration changes proportionally to the osmotic pressure (Fig 2f), which is fully consistent with our finding that cell volume changes proportionally to the osmotic pressure (Fig 2a). Extracting the concentration from the RI and knowing the average cell volumes allows for calculation of the dry mass for each osmotic condition (Fig 2g). The average dry mass of single HeLa cells was 305 +/- 98 pg (Supp Fig 3e) and, as expected, is constant throughout all osmotic shocks (Fig 2g). To directly measure changes of concentration for a single protein within the cytoplasm, we measured the relative change of fluorescence intensity of cells overexpressing cytosolic GFP over time. It also varied proportionally to the osmotic pressure (Supplementary information, Supp Fig 3f-h). Thus, no significant amounts of intracellular solutes were exchanged with the environment, in agreement with previous studies^30, 35^. Our measurement of the dry mass (Fig 2g) also enabled an estimation of the V_OI_. Multiplying the specific volume of dried proteins (0.73 ml/g^54^) by the dry mass, we found *V*_OI_ = 223.34 ± 71.88*μm*3 in agreement with our PBVH fit.

To understand the origin of the rapid recovery after the hypotonic shock, we studied the contributions of various cellular processes involved in the osmotic response, starting with the cytoskeleton. We first imaged the dynamics of the actin cortex during osmotic shocks using SiR-Actin. Upon hypotonic shock, we observed cell blebbing concomitant with cortical actin depolymerization (Fig 3a). Blebs then extended and merged into a large membrane dome (Fig 3a side view). By quantifying cortical actin fluorescence, we observed a complete repolymerization of the cortex four minutes after the shock (Fig 3b), to a value higher than the initial value. Following a hypertonic shock, the actin cytoskeleton appeared more condensed, and its fluorescence intensity gradually increased with time (Fig 3b). We also addressed the behavior of microtubules using SiR-Tubulin. After hypotonic shocks, microtubules also depolymerized and appeared more condensed after a hypertonic shock, but to a smaller extent than actin (Fig 3c-d). These results suggest that actin dynamics is strongly disturbed shortly after osmotic shocks but counteracts high pressure differences by polymerizing over longer times.

**Figure 3.**
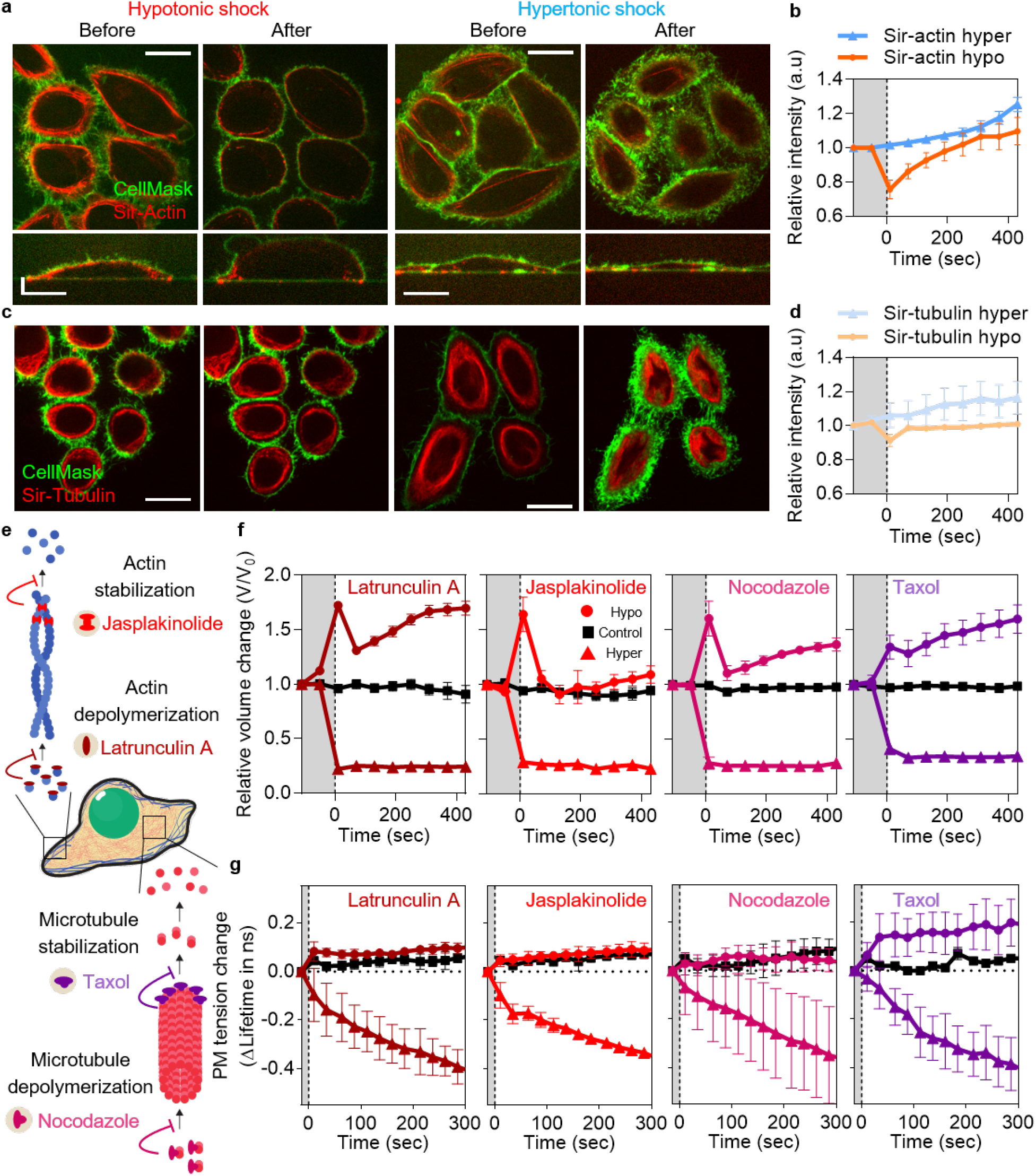
Cytoskeleton controls the long-term response of cells to osmotic shocks. **a**, Actin imaging under hypotonic shock (left) and hypertonic shock (right). Scale bar: 20 *μ*m. Bottom panels are y projections. **b**, Quantification of actin fluorescence intensities during shocks. **c**, Tubulin imaging under hypotonic shock (left) and hypertonic shock (right). Scale bar: 20 *μ*m. **d**, Quantification of tubulin fluorescence intensities during shocks. **e**, Illustrations of cytoskeletal drugs effects. **f**, Single cell volume dynamics of cells treated with latrunculin A, jasplakinolide, nocodazole or taxol during hypotonic shocks (90 mOsm - 75% water, circle), isotonic condition (315 mOsm, square) and hypertonic shocks (700 mOsm - P/P0 = 2, triangle). **g**, Membrane tension dynamics of cells treated with latrunculin A, jasplakinolide, nocodazole or taxol during the same shocks as in e.

To test this hypothesis, we used latrunculin A to depolymerize the F-actin or jasplakinolide to stabilize it (Fig 3e). We then followed the cell volume and tension changes with time and compared them to untreated cells. As described below, none of the drugs used affected the response to hypertonic shocks (Fig 3f, Supp Fig 4a-b), consistent with the hypertonic response being essentially passive. Similarly, both drugs had little effect on the initial peak in cell volume after hypotonic shock, consistent with the short-term response to hypotonicity being passive. However, latrunculin radically modified the later-time recovery compared to non-treated and jasplakinolide-treated cells. Indeed, the volume of latrunculin-treated cells partially recovered after the initial peak, but then diverged a few minutes after shock (Fig 3g). By contrast, the volume of jasplakinolide treated cells evolved similarly to that of non-treated cells (Fig 3f), although over a shorter time scale. Interestingly, the tension dynamics of both latrunculin and jasplakinolide-treated cells were completely decoupled from volume dynamics, as no peak, and thus no recovery, was observed (Fig 3g). Thus, the actin cortex is a major component of the coupling between membrane tension and volume dynamics (Fig 3g).

**Figure 4.**
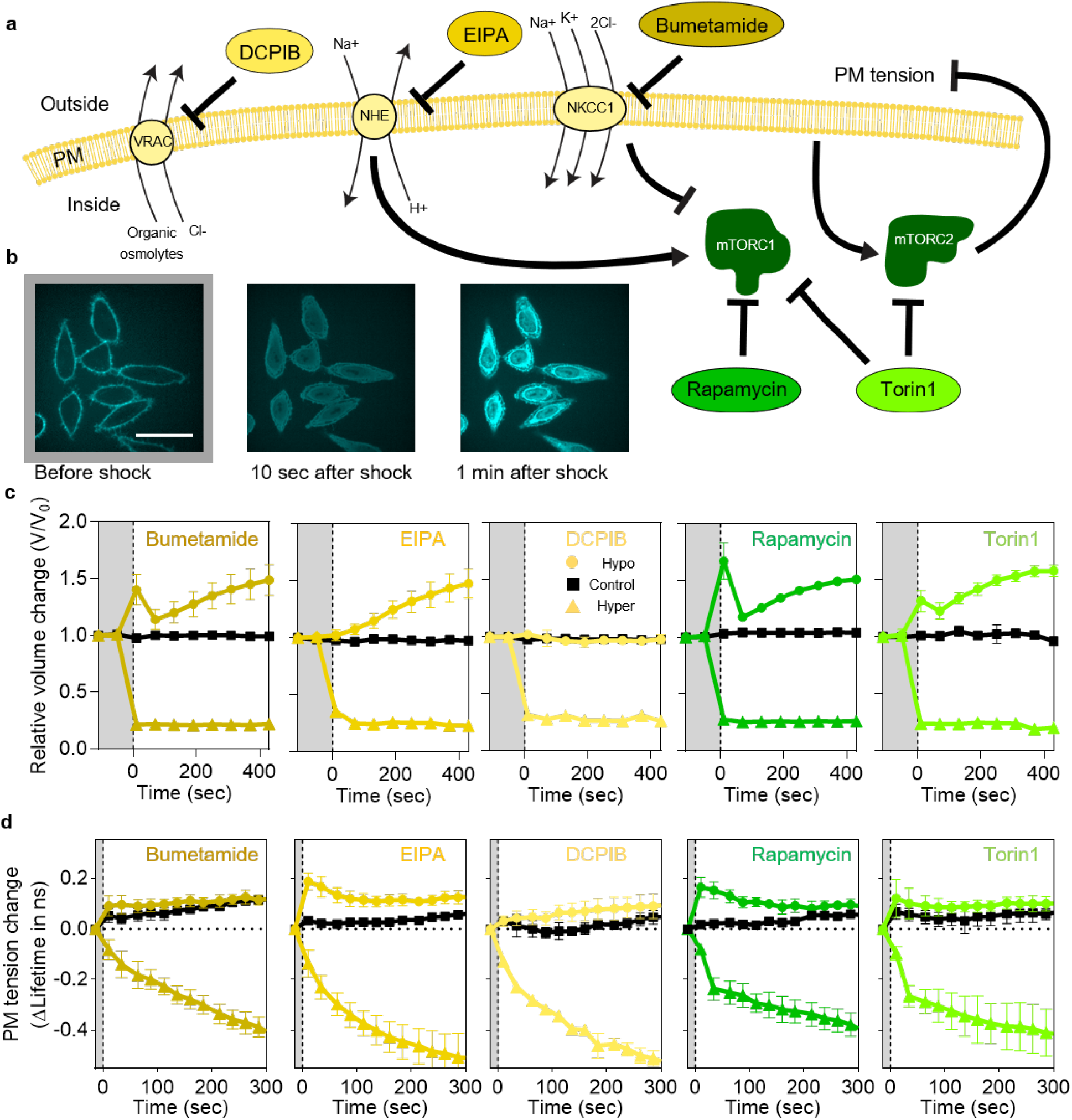
Ion channels are responsible for the short-term response of cells to osmotic shocks. **a**, Illustrations of DCPIB, EIPA and Bumetamide pharmacological effects on, respectively, VRACs, NHE and NKCC1 channels. Signaling pathways from channels to mTOR complexes inhibited by Torin 1 (mTORC1 and mTORC2) or rapamycin (mTORC1) are represented. **b**, Confocal images of DCPIB treated cells and response under hypotonic shock. Scale bar = 40*μ*m. **c**, Single cell volume dynamics in cells treated with Bumetamide, EIPA, DCPIB, rapamycin or Torin1 for hypotonic shocks (90 mOsm - 75% water, circle), isotonic condition (315 mOsm, square) and hypertonic shocks (700 mOsm - P/P0 = 2, triangle). **d**, Membrane tension dynamics in cells treated with Bumetamide, EIPA, DCPIB, rapamycin or Torin1 for identical shocks as in c.

Depolymerization of microtubules with nocodazole had limited effects on the volume dynamics after hypotonic shocks but also decoupled tension from volume changes, as no tension changes were observed (Fig 3f, Supp Fig 4b). Conversely, stabilizing microtubules with taxol clearly affected the dynamics of volume changes, as its peak was significantly smaller than in non-treated cells, and no recovery was observed. In taxol-treated cells, tension dynamics closely followed cell volume dynamics even though they were different from non-treated cells (Fig 3g). These results show that tension dynamics can be decoupled from volume dynamics when actin turnover is affected. As seen for taxol-treated cells, it is possible to qualitatively change the volume and tension response to osmotic shock, while preserving their coupling. Finally, none of the treatments affected the hypertonic response, suggesting that cells respond passively to this condition, or at least without the involvement of the cytoskeleton.

Ion channels are a central regulators of cell volume, and are thus involved in the cell response to osmotic shocks. We used pharmacological inhibitors of the three classes of channels involved in osmotic stress response: DCPIB inhibits VRACs, while EIPA inhibits NHE channels and Bumetanide inhibits NKCC1 channels (Fig 4a). In all hypertonic conditions, none of the inhibitors tested significantly affected the cell volume and tension responses, again indicating that the hypertonic response is essentially passive (Supp Fig 4c). However, after hypotonic shocks, we observed a gradual impact of drugs from Bumetamide to DCPIB on the short-term cell swelling. DCPIB-treated cells were instantaneously permeabilized upon strong hypotonic shock, as seen by the instantaneous labelling of intracellular membranes with CellMask (75% water/90mOsm, Fig 4b). Cell volume did not change in milder hypotonic conditions (25% and 50% water, Fig 4c). By contrast, cells treated with Bumetamide had a smaller but significant peak in cell volume (Fig 4c), and EIPA-treated cells showed no peak immediately after hypotonic shock (Fig 4c). In EIPA- and Bumetamide-treated cells, cell volume slowly diverged three minutes after shock (Fig 4c), while DCPIB-treated cells showed no volume change (Fig 4c). For all three drugs, the tension dynamics during hypotonic shock was strongly affected (Fig 4d). For Bumetamide and EIPA, the response was clearly decoupled from the volume change, while tension remained constant for DCPIB-treated cells, perfectly matching the volume dynamics. Overall, these results show that ion channels that participate in the osmotic balance of the cell, also participate in the coupling between tension and volume changes during osmotic shocks.

The rapid recovery of cell volume and tension during hypotonic shocks shows that these parameters are tightly and actively regulated by the cell. mTORC1 and mTORC2 are involved in cell volume regulation^55^ and membrane tension regulation^29^. Both mTORC1 and mTORC2 are organized around the kinase mTOR, whose phosphorylation can be inhibited using Torin1 which blocks both complexes^56^ while rapamycin is a specific inhibitor of mTORC1^57^. Knowing that both TORC1 and TORC2 are inhibited by hypertonic shocks^24 26, 58^ while TORC2 and mTORC2 are activated by hypotonic shock^26, 28, 29^, we studied the effect of Torin1 and rapamycin treatments on the cell response to osmotic shocks. Volume changes induced by hypotonic shocks were only mildly affected by rapamycin, while Torin1-treated cells exhibited a reduced volume peak after hypotonic shocks (Fig 4c) when compared to non-treated cells. This suggests a more central role of mTORC2 in comparison to mTORC1 in controlling volume and tension response. To test this further, we looked at the dynamics of the tension in both rapamycin and Torin1-treated cells. We observed an initial peak of tension in rapamycin-treated cells similar to non-treated cells, followed by a slower recovery of tension. In Torin1-treated cells, no tension changes were observed (Fig 4d). These results suggest that mTORC2 controls the initial volume/tension coupling, while mTORC2 and mTORC1 are involved in the long-term recovery of both volume and tension. Both rapamycin and Torin1-treated cells did show significant changes of their volume and tension responses to hypertonic shock, strongly supporting that the cell response to hypertonic shock is essentially passive (Supp Fig 4d).

## Discussion

Our study highlights the quantitative relation between cell volume changes and cell tension changes. We showed that cell volume changes are mainly due to cytoplasmic volume changes as opposed to changes in the nucleus volume and confirms that cells modulate their volume according to the PBVH relation. Interestingly, we observed two phases of cell volume response to osmotic shocks: the short-term response - a few seconds after the shock - which was characterized by cell volume variations according to the PBVH relation. The second phase - a few tens of seconds to minutes after the shock - which we called the long-term response, and which was characterized by an asymmetric recovery. Indeed, cell volume recovered fast from hypotonic shocks, but did not recover from hypertonic shocks. This asymmetry of response is probably linked to active counteracting cell processes aimed at mitigating the immediate threat to cell life by increased membrane tension.

During those two phases, we observed that membrane tension followed cell volume changes. In the short-term response, evolution of tension with volume changes was consistent with a model based on membrane unfolding. Fits to the model yielded an estimate of the size of membrane ruffles and invaginations – 37nm – consistent with the smallest membrane structures described in the literature, suggesting that these structures are responsible for the majority of the mechanical response. It also enabled inferring the change of tension according to the change of pressure applied outside the cells. This result is qualitatively maintained during the long-term response, as tension dynamically evolves with the same asymmetry as volume after hypotonic and hypertonic shocks. These results establish that tension passively follows volume changes during the entire duration of the response and recovery to osmotic shocks.

However, this passive coupling between the membrane tension and the cell volume was regulated by active processes of the cell such as the cytoskeleton, the ion channels and mTOR pathways. By disrupting actin or depolymerizing microtubules, we observed no difference in the short-term response of cell volume, consistent with the PBVH relation which does not account for the role of the cytoskeleton. In the long-term response to a hypotonic shock, membrane tension did not vary at all while cell volume fully recovered or increased after the recovery, implying a disruption of the coupling between volume and tension. Conversely, when microtubules were stabilized, the coupling between tension and volume change was preserved, even if the overall response was dramatically changed. We also found that blocking ion channels strongly interfered with the cell volume and tension response but their coupling was only affected for the channels transporting sodium (NHE and NKCC1) as opposed to VRACs. Inhibitors of the mTOR pathways strongly decoupled tension and volume responses on the long term response but only the inhibition of mTORC2 lead to a decoupling on the short term response supporting the notion that mTOR signaling is required for adapting the tension to volume changes. Altogether, those result support the hypothesis of a regulation of the volume and the tension independent from the regulation of their coupling.

Overall, our results support the notion that a large excess of membrane is stored into ruffles maintained by the cytoskeleton, and that the recovery is required to restore this large excess. When the cell volume dramatically increases because of hypotonicity, the cell initially responds by depolymerizing the cytoskeleton to drive membrane unfolding, which results in a release of membrane surface area. The initial volume recovery is mediated through ion channels, as the cytoskeleton is still disrupted, and finalized with actin repolymerization to refold the membrane, under the control of mTOR signaling. Our results show that the coupling between tension and volume is actively regulated by the cytoskeleton, ion channels and mTOR signaling to maintain a quantitative relation between volume and tension well described by passive physical mechanisms.

## Supplementary information

### Nucleus volume change upon osmotic shock

Cells are composed of two main compartments, the cytoplasm and the nucleus. To determine their respective volume changes, we used fluorescence imaging to measure the volume of the nuclei of Hela cells expressing Lamin-B1-GFP – a component of the nucleus membrane – and measure their volume changes under osmotic shocks. The distribution of initial nuclear volumes is narrower than that of overall cell volumes (Supp Fig 1e) centered on 800± 150 *μm*^3^. The qualitative behavior of nuclei under osmotic shocks was similar to that of the whole cells: their volume rapidly increased two-fold after hypotonic shocks and then relaxed back, while they rapidly and stably decreased after hypertonic shocks (Supp Fig 1f). In prior studies^37^, no change of volume was detected under hypotonic shock probably because of low time resolution. Before osmotic shocks, nuclei occupied 24% of the total cell volume. Under hypotonic shocks (75% water), nuclei represented only 17% of the total cell volume while under hypertonic shocks (1M sucrose), nucleus represents 46% of the total cell volume (Supp Fig 1g). Those observations highlight that cell volume changes are essentially due to cytoplasmic volume changes.

### GFP concentration

To directly measure changes of concentration for a single protein within the cytoplasm, we measured the relative change of fluorescence of cells overexpressing cytosolic GFP over time (Supp Fig 3f). Overall, the dynamics of GFP fluorescence was consistent with that of volume and tension: a fast decrease in fluorescence (corresponding to volume increase) followed by a fast recovery after hypotonic shocks (Supp Fig 3g), as opposed to a rapid and stable increase of fluorescence after hypertonic shocks (Supp Fig 3g). As expected, the variation of GFP intensity was always inversely proportional to the volume changes (Supp Fig 3h).

## Methods and Materials

### Cell culture

Human cervical adenocarcinoma cells HeLa-Kyoto were cultured in DMEM, 4.5g/L glucose (61965-026, Thermofischer) supplemented with 10% Fetal Bovine Serum FBS (10270-106, Thermofischer) and 1% Penicillin Streptomycin (P/S, Life Technologies, Carlsbad, CA, USA) in a 5% CO2 incubator (Thermo Scientific, Waltham, MA, USA). HeLa Kyoto EGFP-LaminB1/H2B-mCherry cells from cell lines service (CLS, 330919) were used to image the nuclear membrane. Selection pressure for the stably expressed constructs was kept by adding 0.5mg/mL of G418 and 0.5ug/mL of puromycin in otherwise identical medium as described above. HL-60/S4 cells (ATCC Cat CRL-3306) were cultured in RPMI medium (ATCC-modification) supplemented with 10% FBS and 1% P/S in a 5% CO2 incubator. For all cell lines, number of passages was kept under 20. Our cells were authenticated by Microsynth and are mycoplasma-negative, as tested by GATC Biotech and are not on the list of commonly misidentified cell lines maintained by the International Cell Line Authentication Committee.

### Apply osmotic shock for live cell imaging

Cells were seeded into 35-mm MatTek glass-bottom microwell dishes and were imaged in Leibovitz medium (Thermofischer, 21083027) supplemented with 10% Fetal Bovine Serum and 1% Penicillin Streptomycin. For hypotonic shocks, we simply diluted the cell imaging media with 25%, 50% or 75% of MilliQ water (285 mOsm, 200 mOsm, 125 mOsm). For hypertonic and isotonic shocks, a stock solution of Leibovitz and sucrose (2M) was diluted to obtain a final osmotic pressure of 550 mOsm, 900 mOsm, 1300 mOsm, 2000 mOsm and 3500 mOsm as well as additional intermediate solutions (Supp Fig 1a-b). We chose sucrose over salts to avoid changing specific ion concentrations in the medium. Also, sucrose does not cross the cell membrane, and is not metabolized by HeLa cells. Control hypertonic shocks performed using sorbitol gave identical results (Supp Fig 5a-b). Under the microscope, 1 mL of shock solution was added (1-2s) to the imaging dish containing 1mL of isotonic buffer during imaging. Osmotic shocks were applied 10 seconds before the third time point (2 min). At the end of each experiment, the remaining buffer was collected and its osmotic pressure was measured using an osmometer (Camlab). Since osmotic shocks are applied lived under the microscope, no mixing of the solutions is possible. Using fluorescein, we measured the dilution factor by fluorescence comparing the fluorescence image of the fluorescein solution alone in a dish and the fluorescence of the solution around the cells after added it to 1 mL of medium. As sucrose solutions tend not to mix well with other aqueous solutions and to sediment to the bottom of the imaging chamber, osmolarities of final solutions were corrected by this dilution factor to obtain osmolarities to which cells were subjected.

### Image acquisition and analysis for cell volume measurement

Z-stacks were acquired through a spinning-disk confocal composed of a Nikon Ti-E system, a Yokogawa CSU-X1 Confocal Scanner Unit, a iXon camera (Andor, Belfast, NIR, UK), a Laser stack by Intelligent Imaging Innovation Inc (3i, Denver, CO, USA), a 37°C incubator (Life Imaging service, Basel, Switzerland). All images were acquired with Slidebook software (3i, Denver, CO, USA). In order to measure single cell volume changes through time, HeLa Kyoto cells were labelled with the plasma membrane marker CellMask (Thermofischer C10046). Dyes were diluted in cell medium at 1:1000, incubated at 37°C for 10 min. Confocal Z-stacks (400 nm steps) were acquired every minute for 10 minutes. Osmotic shocks were applied 10 seconds before the third timepoint. Cell 3D images were segmented using a Matlab home-written code, validated with the Limeseg plugin^59^ (Supp Fig 5b) and cell volume and area were extracted. Cells were segmented using a 3D watershed with an intensity threshold automatically determined according to the stack pixel distribution of the entire stack. The tracking in time was straightforward since cells are not moving. Code available on https://github.com/ChloeRoffay/3D-segmentation-time-tracking.

### High-throughput 2D imaging of HL-60/S4 cells for volume estimation

HL-60/S4 cells were collected by centrifugation for 5 min at 180g and resuspended in an osmolarity-adjusted measurement buffer (MB). MB was based on Leibovitz’s L15 medium (no. 21083027, Thermo Fisher Scientific) supplemented with 10% heat-inactivated FBS, 1% penicillin–streptavidin and 0.6% (wt/vol) methyl cellulose (4,000 cPs; Alfa Aesar) for increased viscosity that prevents cell sedimentation during the measuremnts. The osmolarity of MB was adjusted by addition of sucrose or by mixing Leibovitz’s/FCS-based MB with water-based MB of same viscosity, and measured before each experiment using a freezing point osmometer (Fiske 210 Micro-Sample Osmometer, Advanced Instruments). 2D bright-field cell images were acquired at high throughput using real-time deformability cytometry^40^ according to previously published procedures^60^. In brief, the cells suspended in MB were introduced to the microfluidic chip with a syringe pump. The total flow rate was set to 0.16 *μ*Ls^−1^ (0.04 *μ*Ls^−1^ sample flow together with 0.12 *μ*Ls^−1^ focusing sheath flow). The time between resuspension of cells and start of the measurement amounted to roughly 2 min, after that few thousand events were recorded every minute to follow the cell volume response over time. Cell images were acquired at the end of a 300-*μ*m-long channel with a 30×30 *μm*^2^ square cross-section at 2,000 frames per second using stroboscopic illumination with a pulse duration <3 *μ*s to avoid motion blurring. The cell contours were detected in real time and cell area and further parameters were estimated online. Acquired events were filtered for area between 50–500 *μm*^2^ to excluded debris and area ratio between 1.00–1.05 to excluded rough or incomplete contours^60^. Volume of the cells was estimated offline by 360° rotation of upper and lower halves of the 2D cell contours around the symmetry axis, and averaging the two obtained values^61^ using ShapeOut version 1.0.1 (available at https://github.com/ZELLMECHANIK-DRESDEN/ShapeOut).

### Tube pulling experiment

Membrane nanotube pulling experiments were performed on the setup published in^62^ allowing simultaneous optical tweezer application, spinning disc confocal and brightfield imaging based on an inverted Nikon eclipse Ti microscope and a 5W 1064nm laser focused through a 100 1.3 NA oil objective (ML5-CW-P-TKS-OTS, Manlight). A membrane nanotube was formed by displacing the cell’s observation chamber away from a concanavalin-A-coated bead (3.05 mm diameter, Spherotec) held the optical trap, and previously in contact with the cell to adhere to the cell membrane. The force F exerted on the bead was calculated from Hooke’s law: F = k.x, where k is the stiffness of the trap (k = 8.58 pN.pix^−1^.W^−1^) and x is the displacement of the bead from its initial zero-force position.

### Image acquisition and analysis for Flipper-TR imaging

Membrane tension measurement were performed on the setup published in^33^. Setup used for imaging is a Nikon Eclipse Ti A1R microscope equipped with a time-correlated single-photon counting module from PicoQuant. Excitation was performed using a pulsed 485 nm laser (PicoQuant, LDH-D-C-485) operating at 20 MHz, and the emission signal was collected through a 600/50 nm bandpass filter using a gated PMA hybrid 40 detector and a TimeHarp 260 PICO board (PicoQuant). In order to measure membrane tension changes through time, HeLa Kyoto cells were labelled with Flipper-TR (Spirochrome SC020). Flipper-TR was dissolved in DMSO at 1 mM stock solutions. Cells were labelled with a 1:1000 dilution from the DMSO stock, incubated 37°C for 15 min and slices were acquired every 25 sec for 10 minutes (Fig 3C) without washing the probe. Osmotic shocks were applied 10 seconds before the second timepoint. Quality of imaging is altered in DMEM (with or without FBS, independently of phenol-red), such that all images were acquired in Leibovitz for short-term imaging (less than 2h) or FluoroBrite (A1896701) for longer times. Lifetimes of Flipper-TR were extracted from FLIM images using SymPhoTime 64 software (PicoQuant) by fitting fluorescence decay data from all pixels to a dual exponential model after deconvoluting the instrument response function (calculated by the software). We selected full images instead of choosing region of interest because the fitting was mildly affected and the result were more reproducible. The full-width at half-maximum response of the instrument was measured at 176 ps.

### Refractive index measurement and processing

The three-dimensional (3D) refractive index (RI) distribution of samples was measured using a custom-made ODT (Optical Diffraction Tomography) microscope. The optical setup of ODT employs Mach-Zehnder interferometry in order to measure complex optical fields of light scattered by samples from various incident angles, as shown in^52^. A coherent laser beam (wavelength = 532 nm, frequency-doubled Nd-YAG laser, Torus, Laser Quantum, Inc., UK) is divided into two beams by a 2×2 single-mode fiber optic coupler. One beam is used as a reference beam and the other beam illuminates the specimen on the stage of a custom-made inverted microscope through a tube lens (f = 175 mm) and a high numerical aperture (NA) objective lens (NA = 1.2, 63x, water immersion, Carl Zeiss AG, Germany). To reconstruct a 3D RI tomogram of samples in a field-of-view, the samples are illuminated with 150 various incident angles scanned by a dual-axis galvanomirror (GVS012/M, Thorlabs Inc., USA). The diffracted beam from a sample is collected by a high NA objective lens (NA = 1.3, 100x, oil immersion, Carl Zeiss AG) and a tube lens (f = 200 mm). The total magnification is set to be 90.5x. The beam diffracted by the sample interferes with the reference beam at the image plane, and generates a spatially modulated hologram. The hologram is recorded with a CCD camera (FL3-U3-13Y3M-C, FLIR Systems, Inc., USA). From measured holograms, the 3D RI tomograms are reconstructed by the Fourier diffraction theorem employing the first-order Rytov approximation^51, 63^. Cells were manually segmented based on the epifluorescence image of the membrane (CellMask staining as previously described). Cell’s basis was automatically detected to correct the z-drift. By applying the segmented ROI to the projection of the RI tomogram onto the cell’s basis plane, pixels value of RI were extracted, averaged over the entire cell, and averaged over many cells for each osmotic condition.

### Image acquisition and analysis of the cytoskeleton

In order to measure actin or tubulin intensity changes through time, HeLa Kyoto cells were labelled with the plasma membrane marker Cell Mask orange (Thermofischer C10045) and SiR-actin (SC001, Spirochrome) or SiR-tubulin (SC002, Spirochrome). Dyes were dissolved in DMSO at a stock concentration of 1mM, and diluted 1:1000 in cell’s medium, incubated at 37°C for 20 min to label cells. Cells were then imaged as described above. Cells were manually segmented and average intensity was extracted using ImageJ.

### Drug treatment

Concentrations of drugs was kept constant through the experiment. Cell were initially incubated in culture DMEM with drugs at 37°C (see below drugs concentration, incubation time). DMEM was replaced by Leibovitz with drugs and CellMask or Flipper-TR. The osmotic shocks were applied as before except that the solution contains the same drugs concentration. The following pharmacological inhibitors and chemical compounds were used: 50 nM Latrunculin A for 1h (SIGMA L5163), 200 nM Jasplakinolide for 30 min (ENZO ALX-350-275), 5 *μ*M Nocodazole for 30 min (SIGMA M1404), 1 *μ*M Taxol for 1h (SIGMA T1912), 100 *μ*M DCPIB for 30 min (TOCRIS 1540), 50 *μ*M EIPA for 30 min (TOCRIS 3378), 100 *μ*M Bumetamide for 30 min (SIGMA B3023), 250 nM Torin1 for 30 min (LC Lab T-7887) and 100 nM Rapamycin for 30 min (LC Lab R-5000).

### GFP experiments

pGFP was transfected using FuGENE^®^6 Transfection Reagent (E2691,Promega) with OptiMEM (31985088, Thermofischer). Transfected Hela Kyoto cells were image with the same spinning disk confocal set-up, single confocal planes being acquired every second for 10 minutes. Nikon’s Perfect Focusing System was used to keep focus during the entire osmotic shock. Cells were manually segmented using ImageJ and the mean fluorescence was extracted through time. For each cell, values were normalized to the initial value.

All images were constructed using ImageJ. All graphs were constructed with GraphPad Prism 8.

## Acknowledgments

We thank K. Roux, R. Wimbish and N. Kléna for critical reading of the manuscript, and C. Tomba for discussions. AR acknowledges funding from Human Frontier Science Program Young Investigator Grant RGY0076/2009-C, the Swiss National Fund for Research Grants N31003A_149975 and N31003A_173087, and Synergia Grant N CRSII5_189996, the European Research Council Consolidator Grant N 311536, and Synergy Grant N951324-R2-TENSION.

## Author contributions

C.R. and A.R. designed the research. C.R., G.M., K.K., V.B., M.U. performed all experiments and image analyses. V.M., J.GC., S.M, J.G. contributed with tools and technique. M.L. did the theoretical models. C.T. and A.R. wrote the manuscript, with editions from K.K, M.U., V.M., J.G and M.L.

---

**Figure 1. Supplementary.**
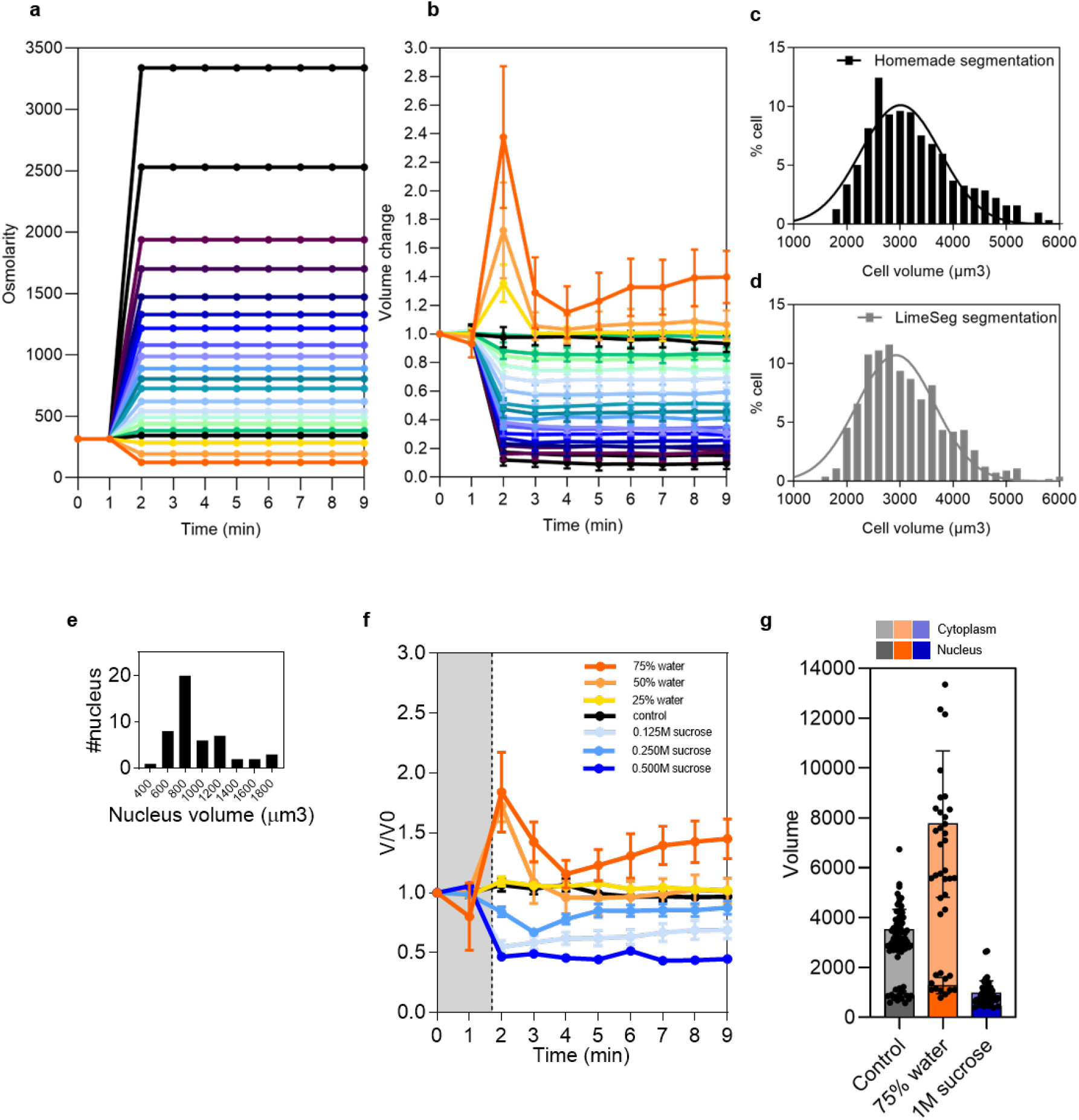
Measure cell volume, nucleus volume and osmotic pressure. **a**, Osmolarity changes during time. **b**, Volume changes during time. **c**, Cell volume distribution in isotonic conditions measured with the homemade segmentation (n = 959). **d**, Cell volume distribution in isotonic conditions measured with the Limeseg segmentation (n = 578). **e**, Distribution of nucleus size in isotonic medium. **f**, Single nucleus volume dynamic under osmotic shock. **g**, Relative contribution of nucleus and cytoplasm to cell volume changes under hypotonic shock (75% water - orange) and hypertonic shock (1M sucrose - blue).

**Figure 2. Supplementary.**
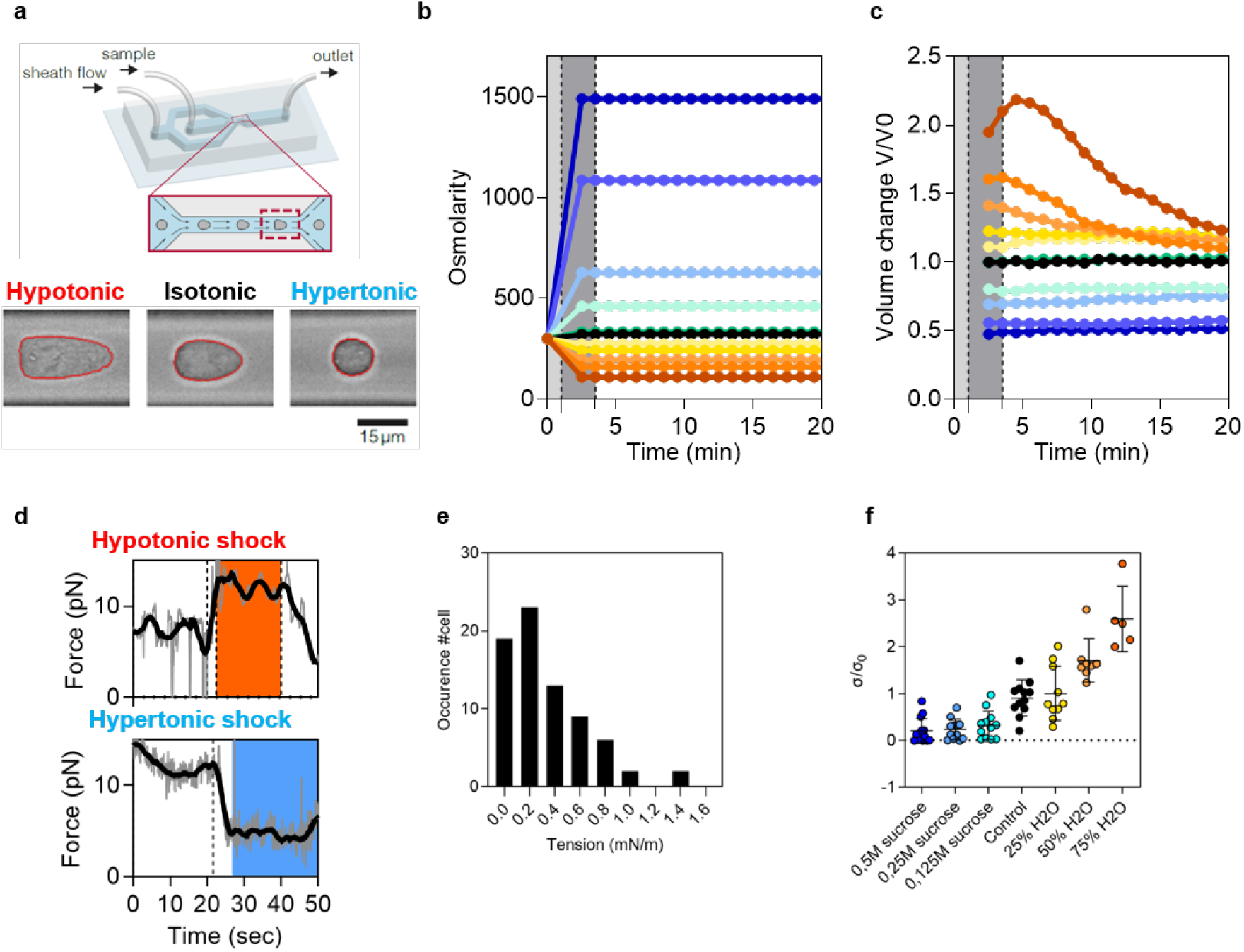
Measure nucleus volume, use RT-DC and measure tension with tube pulling. **a**, RT-DC principles. **b**, Osmolarity changes under osmotic shocks with RT-DC measurement. **c**, Cell volume dynamic under osmotic shocks with RT-DC measurement. **d**, Force measurement under osmotic conditions (top hypotonic shock, bottom hypertonic shock). **e**, Distribution of initial tether force to pull a cell membrane tube (isotonic condition). **f**, Relative tension measurement under osmotic conditions.

**Figure 3. Supplementary.**
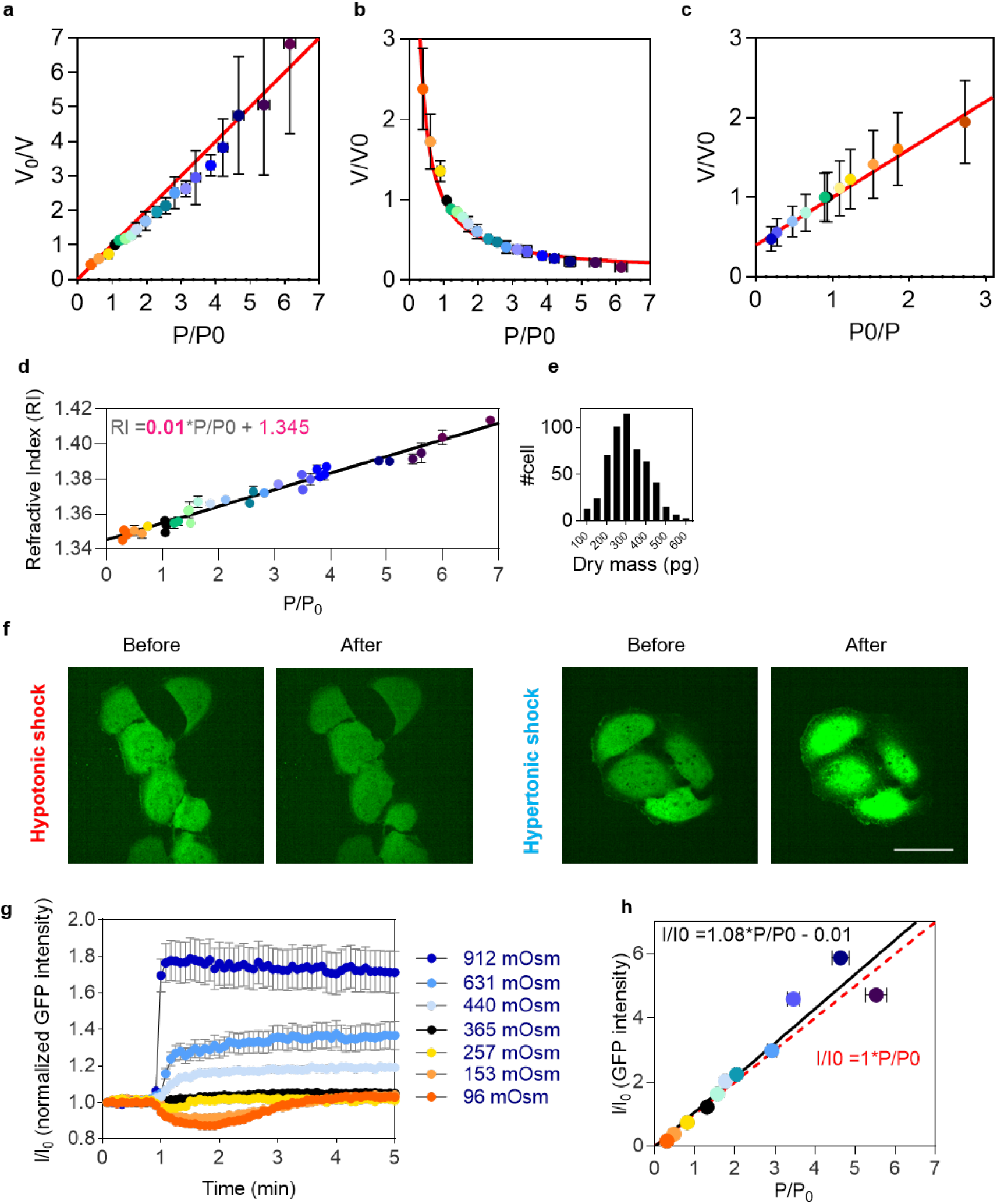
Quantitative coupling of cell membrane tension to osmotic shocks and diffusion data. **a**, Inverse of volume change (V0/V) according to pressure change (P/P0) in HeLa cells. **b**, Volume change (V/V0) according to pressure change (P/P0) in HeLa cells. **c**, Ponder’s relation measuring volume with RT-DC in HL-60/S4 cells. **d**, Refractive index of cell according to change of pressure (P/P0). **e**, Distribution of dry mass (pg) in isotonic medium. **f**, GFP tranfected cells under hypotonic shock (left) and hypertonic shock (right). **g**, Relative change of intensity (I/I0) in time for various osmolarities. **h**, Relative change of intensity (I/I0) according to pressure changes (P/P0).

**Figure 4. Supplementary.**
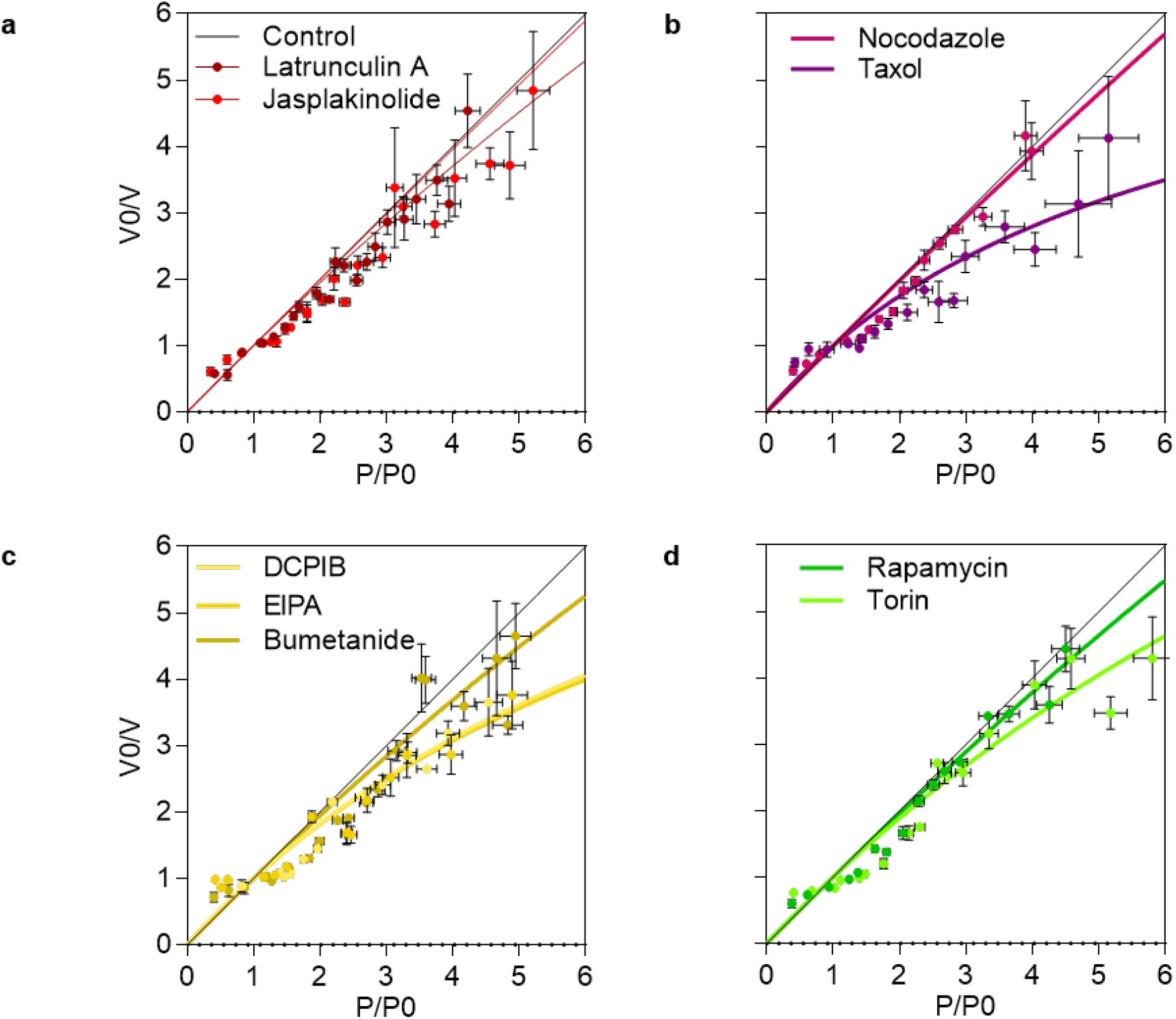
Ponder’s relation and cell volume under drug treatment. **a**, Ponder’s relation for latrunculin A or jasplakinolide treated cells. **b**, Ponder’s relation for nocodazole or taxol treated cells. **c**, Ponder’s relation for DCPIB or EIPA or Bumetamide treated cells. **d**, Ponder’s relation for rapamycin or Torin1 treated cells.

**Figure 5. Supplementary.**
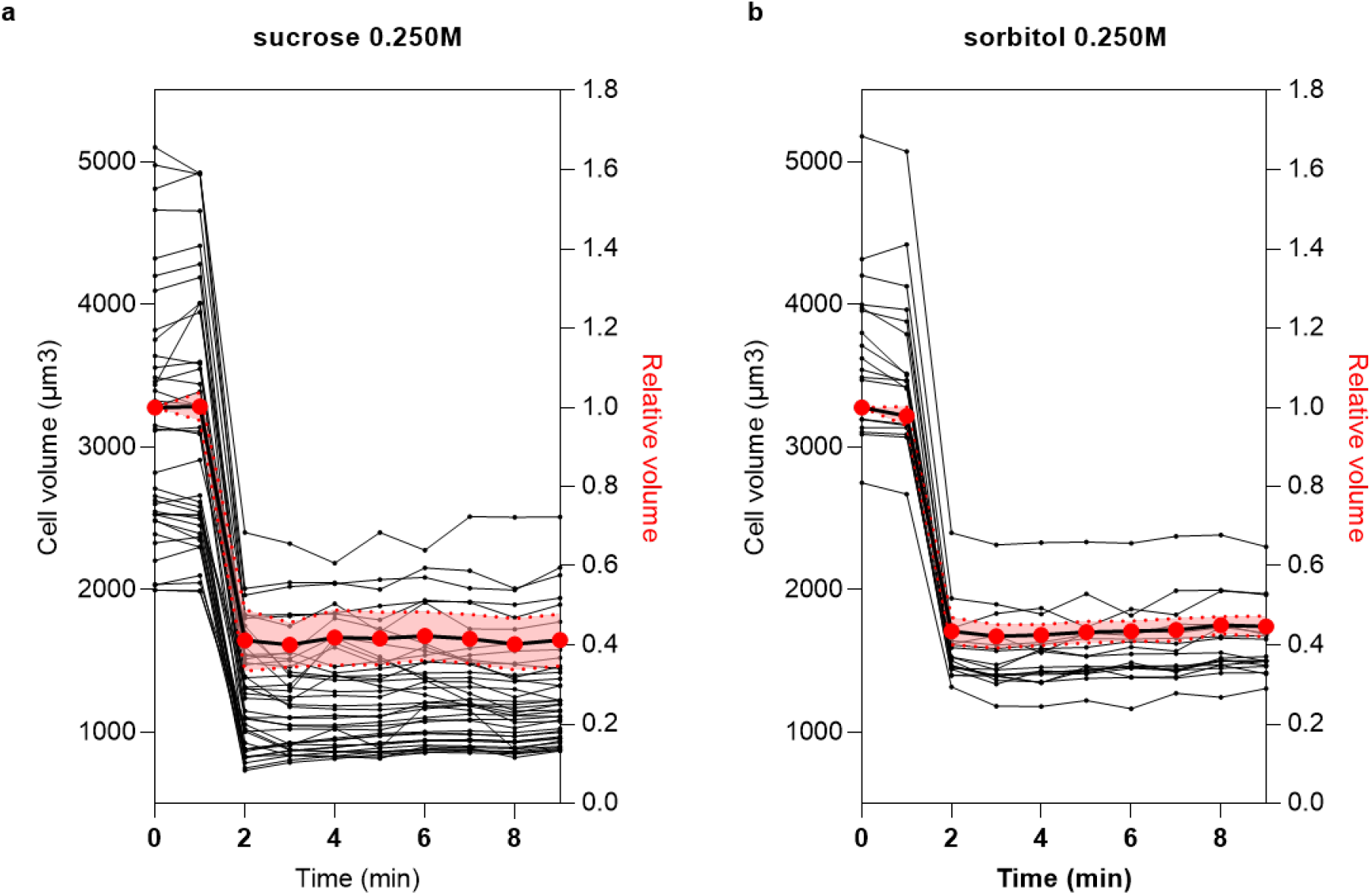
Protein concentration changes using refractive index (RI) measurement. **a**, Absolute volume of cells undergoing a sucrose hypertonic shocks 0.250M (each black line are individual cells) and relative volume changes of all the cells. **b**, Absolute volume of cells undergoing a sorbitol hypertonic shocks 0.250M (each black line are individual cells) and relative volume changes of all the cells.

## Supplemental mathematical modeling

To account for the experimentally observed relationship between membrane area and tension shown in Fig. 2b of the main text, we assume that a tensionless membrane is not flat. Instead, its curvature is dictated by a collection of protein scaffolds of possibly disparate types, including caveolins, other membrane-bound proteins and attachments to the cytoskeleton. As the membrane tension is increased, these “ruffles” unfold, increasing the membrane’s projected area. Our model is athermal, as the additional excess membrane area stored in the membrane’s thermal fluctuations is in practice small compared to the areas considered here.

The Hamiltonian for the roughened membrane reads

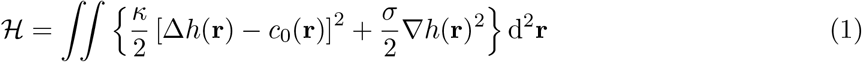

where *h*(**r**) is the height function of the membrane. This is the standard Helfrich Hamiltonian in the Monge gauge, except for a spacially varying spontaneous curvature *c*_0_(**r**) representing the built-in curvature due to membrane-protein interactions. We define *c*_0_(**r**) as a random function, reflecting the messy character of the static protein-induced membrane ruffles. Specifically, we make it a Gaussian variable with a characteristic correlation length *a* that fixes the typical size of the ruffles. Formally, this reads

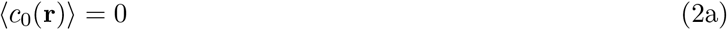

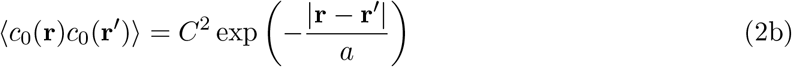

The first equation implies a zero mean curvature. This is not a critical assumption however, as setting a non-zero mean curvature does not change the final result of our calculation. The constant *C* gives the typical magnitude of the local curvature. Fourier transforming the two-point correlator yields

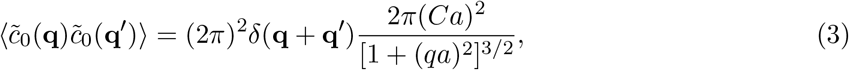

where *q* is the modulus of the wavevector **q**.

Minimizing the Hamiltonian with respect to the height function *h*(**r**) yields the mechanical equilibrium equations for this system. In Fourier space:

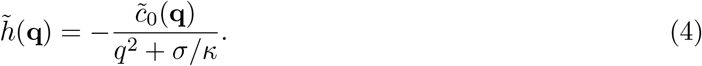

We compute the ratio of the real membrane area to the apparent (projected) area in the Monge gauge as

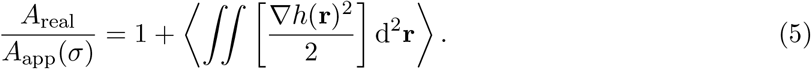

Taking this expression to Fourier space and combining Eqs. (3) and (4) finally yields

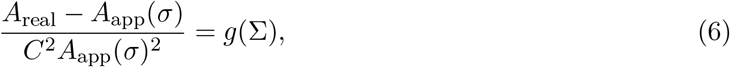

where Σ = *σα*^2^/*κ* and the function *g* is given by

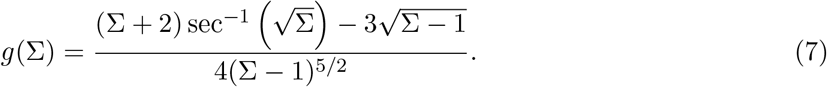

To find the apparent area of the membrane, we solve Eq. (6) for *A*_app_. In the thermodynamic limit, the stored area of the membrane in the zero-tension limit is huge; in other words *C*^2^*A*_real_ → +∞. In this limit the area ratio plotted in Fig. 2b simply reads

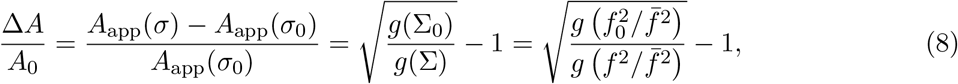

where *σ*_0_ is some reference tension (the isotonic value) and Σ_0_ is its dimensionless counterpart. In the last equality we moreover introduce the directly measurable tube force 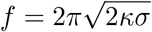, its isotonic value *f*_0_ as well as the characteristic force 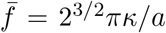. Using the last expression of Eq. (8) as a fitting function for the experimentally measured Δ*A/A*_0_ *vs. f*^2^ curve (where *f*^2^ is measured post-osmotic shock) we find the following values for our fit parameters:

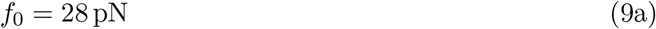

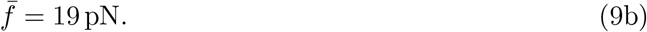

The fit is presented as a line in Fig. 2b. Assuming a bending modulus *κ* = 20*k_B_T* for the membrane inside the tube, these values yield

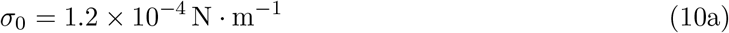

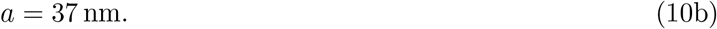

The former value appears pretty typical for a cell membrane tension. The latter value is more informative: the typical size of the membrane ruffles is of the order of a few tens of nanometers.

One might finally ask what fraction of the total area is stored in ruffles under isotonic conditions. According to the model this quantity reads

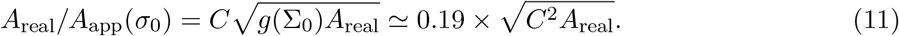

Since the fit does not specify the value of *C*^2^*A*_real_ (it only assumes it is significantly larger than one), it cannot answer this question quantitatively. Conversely, if we assume *A*_real_/*A*_app_(*σ*_0_) = 2.5 and use the experimental value *A*_app_(*σ*_0_) ≃ 775 *μ*m^2^, we get

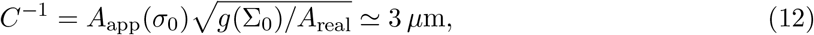

which makes for fairly shallow ruffles, and thus validating our use of the Monge gauge and small-slope expansion of the membrane Hamiltonian.

